# Sex-Specific Modulation of Gene Expression by 17beta-Estradiol in Human Meniscal Cells: Pathways to Targeted Osteoarthritis Therapies

**DOI:** 10.64898/2026.07.24.740612

**Authors:** Youngmin Yu, John Vergis, Hunter M. Eby, Mario Markho, Jonathan Kopacz, Kayla Cartwright, Michael Hershey, Jiayong Liu, Robert McCullumsmith

## Abstract

Osteoarthritis (OA) disproportionately affects women, and estrogen has been implicated in cartilage and joint homeostasis, yet its effects on meniscal fibrochondrocytes (MFCs), and whether those effects differ by sex, remain poorly defined. We reanalyzed a publicly available RNA-sequencing dataset (Gene Expression Omnibus, GSE199087) comprising human MFCs from a male and a female donor treated with 17β-estradiol (E2) or vehicle. Differential expression, Gene Set Enrichment Analysis, Enrichr, and iLINCS were integrated to identify the transcriptional programs modulated by E2 in each sex. In female MFCs, E2 upregulated pathways governing DNA replication, cell-cycle progression, and genomic maintenance (e.g., MCM8, BRCA1, RAD51), while downregulating inflammatory and extracellular matrix-degrading genes, including MMP1, IL12A, CXCL12, and MYD88. In male MFCs, E2 instead upregulated chromatin remodeling and developmental signaling programs, led by SRCAP, ERCC6, and NOTCH1, and downregulated antigen presentation and mitochondrial genes. These divergent responses indicate that E2 engages distinct, sex-specific transcriptional programs in MFCs, with the female profile favoring proliferation and matrix preservation. Although derived from a limited sample and therefore hypothesis-generating, these findings nominate candidate mechanisms underlying sex differences in meniscal biology and OA susceptibility, and warrant validation in larger, sex-balanced cohorts.

**Graphical Abstract:** 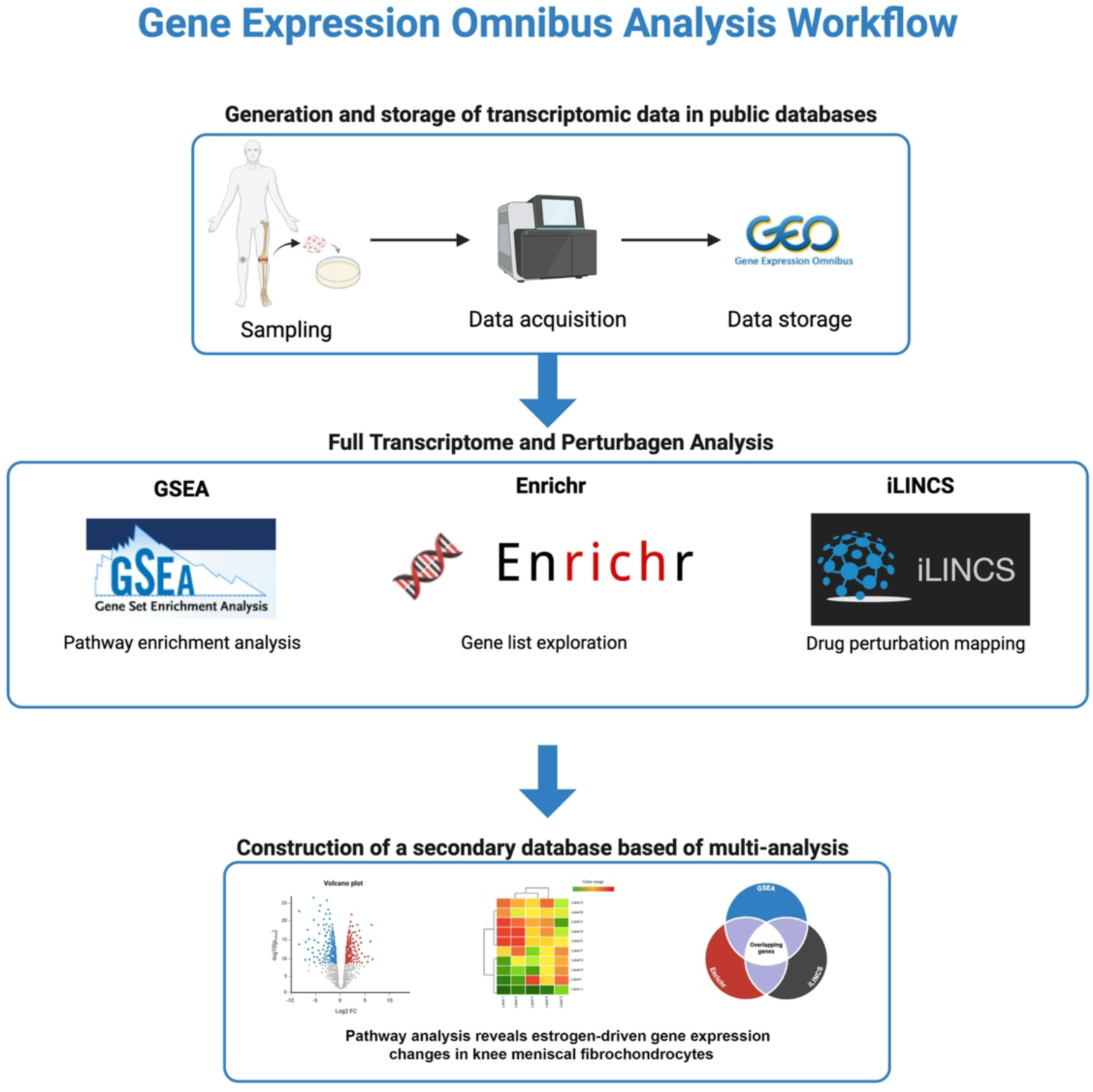

## Introduction

Osteoarthritis (OA) is a debilitating joint disease disproportionately affecting women, especially in post-menopausal years. This disparity suggests a potential role of sex hormones in the pathogenesis and progression of the disease, particularly estrogen (*1*). Research indicates that estrogen influences tissue repair and homeostasis, which are critical in joint health. Notably, estrogen’s interaction with matrix metalloproteinases (MMPs), the principal enzymes driving extracellular matrix (ECM) and collagen breakdown, suggests a complex role in maintaining cartilage integrity (*2, 3*).

Estrogen’s protective effects in cartilage are supported by studies demonstrating its ability to suppress MMP expression, which is crucial in preventing ECM degradation—a hallmark of osteoarthritis. For instance, 17β-estradiol has been shown to reduce the expression of MMPs like MMP-1, MMP-3, and MMP-13 in articular chondrocytes, highlighting its potential chondroprotective effects (*2*). Moreover, the modulation of MMP activity by estrogen may also involve microRNAs, such as miR-140, which further regulates MMP-13, a critical enzyme in cartilage matrix breakdown (*4*).

Despite these protective effects, the role of estrogen in OA is not straightforward. Sex differences in knee biomechanics, such as joint alignment and muscle strength, also contribute to the increased incidence of OA in females compared to males (*1*). These biomechanical factors, combined with the molecular effects of estrogen on cartilage and subchondral bone, underline the complexity of OA pathogenesis in different sexes.

The influence of estrogen on meniscal cells, which play a crucial role in load transmission and shock absorption in the knee, remains underexplored. Meniscal fibrochondrocytes (MFCs) exhibit plasticity that is crucial for the repair of meniscal injuries, and their response to estrogen could provide insights into sex-specific therapeutic targets for OA (*5*). This study aims to delve deeper into the sex-specific effects of 17β-estradiol on MFCs, exploring both the protective and potentially deleterious impacts of estrogen therapy on joint health. By understanding these mechanisms, we can pave the way for personalized and sex-specific treatments for osteoarthritis, which could significantly improve outcomes for patients suffering from this chronic condition (*6*).

## Methods

### Dataset and Bioinformatic Analysis

We reanalyzed RNA sequencing data from the dataset GSE199087, available on the Gene Expression Omnibus (GEO) database. This dataset comprises transcriptomic profiles of human meniscal fibrochondrocytes (MFCs) treated with 17β-estradiol (E2). The samples analyzed in this study were human MFCs collected from the knee of a 47-year-old male cadaveric donor and an age-matched female donor. Cells were collected in three separate preparations, and experiments were conducted in triplicate (treating E2). Cells were treated by pulsed dosing to resemble normal estrogen cycle, where E2 was administered for 1 hour followed by 23 hours of E2-free media. This procedure was repeated three times for 72 hours. Total RNA was extracted and processed using assay-specific kits, followed by sequencing on a NextSeq 550 platform using the NEBNext Ultra II Directional RNA Library Prep Kit.

### Data Processing and Pathway Analysis

For our bioinformatic analysis, we first filtered and aligned the raw reads to the GRCh38 release 101 reference transcriptome using Kallisto. Differential expression analysis was subsequently conducted using Limma-Voom. To ensure robustness and reproducibility, we followed established best practices for reanalyzing published datasets using recent bioinformatics tools (*7–10*). Our data processing rigorously assessed gene expression changes, enabling a detailed exploration of the molecular responses to E2 treatment.

### Volcano Plot

A volcano plot (Figure 1) was generated using GraphPad Prism (version 10.4.1, GraphPad Software, San Diego, CA, USA) to visualize significantly differentially expressed genes (DEGs). The plot represents differential expression as log2 fold-change (log2FC) on the x-axis against the significance (-log10(p-value)) on the y-axis. Genes above the dotted horizontal threshold (-log10(p-value) > 1.7, corresponding to p < 0.02) were considered statistically significant. Previous studies have used a similar significance threshold, typically set at -log10(p-value) of 1.0 or greater (*11–13*). Genes were categorized based on log2 fold-change, with those significantly upregulated or downregulated highlighted distinctly. Selected genes involved in overlapping pathways identified by pathway analyses were labeled directly on the volcano plots.

**Figure 1:**
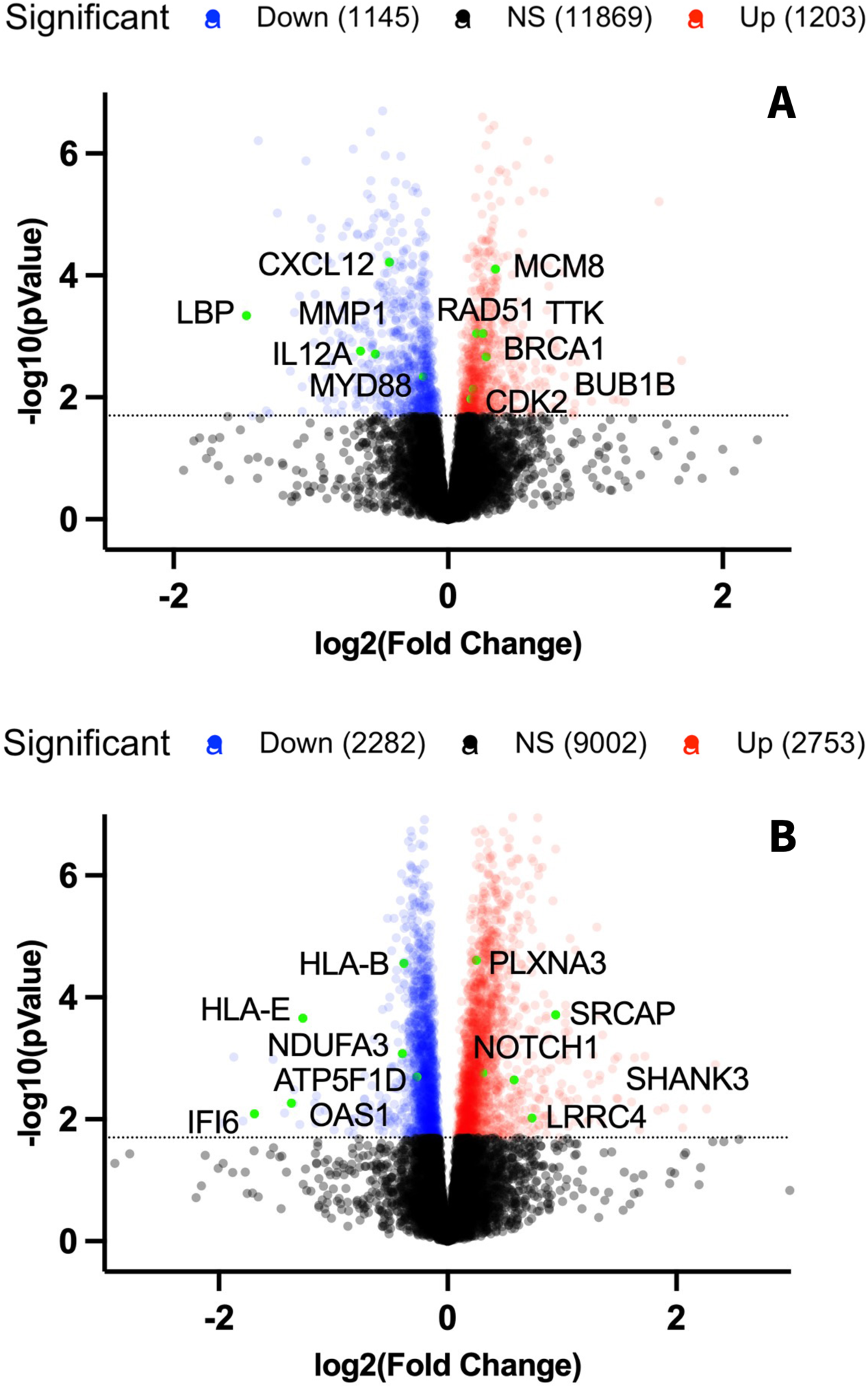
Differential Gene Expression Volcano Plot. (A) Female MFCs E2 treated vs control, (B) Male MFCs E2 treated vs control. -log10(pValue) cut off value of 1.7 (p<0.02) depicted by the dotted horizontal line. Genes involved in overlapping pathways (Table 3, 4) are labeled and plotted in green. MFC = meniscal fibrochondrocytes, E2 = 17β-estradiol

### Overlapping pathway analysis

For pathway analysis, we employed Gene Set Enrichment Analysis (GSEA), Enrichr, and iLINCS to interpret the biological meaning behind the gene expression data. GSEA was used to assess the overrepresentation of gene sets in the ranked list of genes from our dataset, identifying both upregulated and downregulated pathways based on their normalized enrichment scores (NES) and p-values, while Enrichr and iLINCS results are reported as combined scores (CS). We specifically focused on the top overlapping significant pathways identified by GSEA, Enrichr, and iLINCS, considering these pathways as central to our results.

### Leading Edge Gene Analysis

GSEA also provided a leading-edge (LE) gene analysis, which identifies the core subset of genes that contribute most significantly to the enrichment of identified pathways. The Gene Ontology pathway package was used to define gene sets (*14*). This analysis is crucial as it determines the genes that drive the pathway’s association with the condition under study. By analyzing Tags, List, and Signal—metrics that elucidate the contribution of genes to the enrichment score and their position in the ranked list—we were able to pinpoint the genes most critical for the observed changes in pathway activity. The overlap of these leading-edge genes across different significant pathways was further analyzed to identify key drivers of pathway activity and potential targets for therapeutic intervention.

## Results

### Differential Gene Expression Volcano Plot Analysis

#### Female Meniscal Fibrochondrocyte (MFC) Response to 17β-estradiol (E2)

Volcano plot analysis (Figure 1A) of female MFCs following E2 treatment identified significant differential gene expression, highlighting key upregulated genes involved in cell-cycle regulation and DNA repair (e.g., MCM8, RAD51, TTK, BRCA1, BUB1B, CDK2). In contrast, significantly downregulated genes included factors associated with inflammatory and ECM-degrading processes (e.g., CXCL12, LBP, IL12A, MYD88, MMP1).

#### Male Meniscal Fibrochondrocyte (MFC) Response to 17β-estradiol (E2)

The volcano plot for male MFCs (Figure 1B) demonstrated a distinct gene expression response to E2 treatment. Significantly upregulated genes included those associated with synapse assembly, chromatin remodeling, and developmental signaling (e.g., PLXNA3, SRCAP, NOTCH1, SHANK3, LRRC4). Conversely, significantly downregulated genes were linked to immune response and mitochondrial function (e.g., HLA-B, HLA-E, NDUFA3, ATP5F1D, OAS1, IFI6).

### Overlapping pathways in GSEA, Enrichr, and iLINCS

#### Female Meniscal Fibrochondrocyte (MFC) Response to 17β-estradiol (E2)

Analysis of female MFCs revealed significant gene expression shifts following E2 treatment. Differential gene expression analysis identified key upregulated genes involved in cell-cycle regulation and DNA replication, including MCM8, RAD51, TTK, BRCA1, BUB1B, and CDK2. Conversely, genes associated with inflammatory and proteolytic processes such as CXCL12, LBP, IL12A, MYD88, and MMP1 were significantly downregulated (Figure 1A). Pathway analyses combining GSEA, Enrichr, and iLINCS methods further supported these findings. Notably, upregulated pathways included “DNA replication” (GSEA NES = 2.14, Enrichr CS = 70.77, iLINCS CS = 68.19) and “positive regulation of cell cycle” (GSEA NES = 1.78, Enrichr CS = 304.79, iLINCS CS = 37.68) (Table 1A). Downregulated pathways included “endopeptidase activity” (GSEA NES = -1.59, Enrichr CS = -4.45, iLINCS CS = -7.88) and “fibroblast growth factor receptor signaling” (GSEA NES = -1.76, Enrichr CS = -4.09, iLINCS CS = -5.60), aligning with estrogen’s known chondroprotective roles in articular cartilage (Table 1B).

**Table 1:** Female MFCs Overlapping Pathways in GSEA, Enrichr, and iLINCS.

**A**
| GSEA (NES) | Enrichr (CS) | iLINCS (CS) | Pathways |
| --- | --- | --- | --- |
| 2.154360328 | 100.9902121 | 34.39703946 | DNA-templated DNA replication |
| 2.137815011 | 70.77061606 | 68.18824054 | DNA replication |
| 1.808779243 | 48.87510934 | 43.68692806 | DNA strand elongation |
| 1.802502173 | 48.87510934 | 43.68692806 | DNA strand elongation involved in DNA replication |
| 1.799546554 | 80.18720351 | 45.74337059 | mitotic spindle |
| 1.777878983 | 304.7910995 | 37.68206261 | positive regulation of cell cycle process |
| 1.774111895 | 11.9388137 | 34.72680563 | regulation of cell cycle phase transition |
| 1.756572064 | 71.25149797 | 17.04543818 | regulation of mitotic nuclear division |
| 1.725393344 | 60.22313912 | 63.19304242 | spindle |
| 1.68479331 | 11.9388137 | 73.16810303 | mitotic cell cycle checkpoint signaling |

**B**
| GSEA (NES) | Enrichr (CS) | iLINCS (CS) | Pathways |
| --- | --- | --- | --- |
| -1.755192313 | -4.089540345 | -5.59687559 | fibroblast growth factor receptor signaling pathway |
| -1.589677307 | -12.07485637 | -6.180306153 | regulation of vascular associated smooth muscle cell proliferation |
| -1.588787239 | -7.88456186 | -6.508939173 | growth factor receptor binding |
| -1.586038947 | -4.446965571 | -7.875795147 | endopeptidase activity |
| -1.544621658 | -2.66121418 | -6.557556876 | axon guidance |
| -1.544621658 | -1.627177885 | -7.328505222 | neuron projection guidance |
| -1.428664872 | -4.950199324 | -6.060116536 | protein processing |
| -1.421994787 | -1.352608914 | -12.46242425 | axonogenesis |
| -1.322737516 | -3.084080285 | -8.976075593 | regulation of defense response |
| -1.291330917 | -2.383454485 | -6.377318288 | positive regulation of cell death |

Female meniscal fibrochondrocytes overlapping gene expression pathways in GSEA, Enrichr, and iLINCS. (A) shows upregulated pathways. (B) shows downregulated pathways. NES = Normalized Enrichment Score, CS = Combined Score, MFC = meniscal fibrochondrocytes

### Male Meniscal Fibrochondrocyte (MFC) Response to 17β-estradiol (E2)

In male MFCs, distinct sex-dependent responses were observed following E2 treatment. Differentially expressed genes indicated upregulation of factors traditionally associated with synapse assembly, chromatin remodeling, and developmental signaling, such as PLXNA3, SRCAP, NOTCH1, SHANK3, and LRRC4. Concurrently, there was notable downregulation of genes involved in immune response and peptidase activity, including HLA-B, HLA-E, OAS1, IFI6, CST3, and SERPING1 (Figure 1B). Integrated analyses highlighted pathways such as “synapse assembly” (GSEA NES = 1.98, Enrichr CS = 10.35, iLINCS CS = 18.49) and “chromatin remodeling” (GSEA NES = 1.77, Enrichr CS = 6.62, iLINCS CS = 23.60) (Table 2A). Downregulated pathways included “antigen processing via MHC class I” (GSEA NES = -2.07, Enrichr CS = -10.12, iLINCS CS = -63.35) and “regulation of peptidase activity” (GSEA NES = - 1.55, Enrichr CS = -6.49, iLINCS CS = -14.16), suggesting altered immune and proteolytic responses unique to male cells (Table 2B).

**Table 2:**
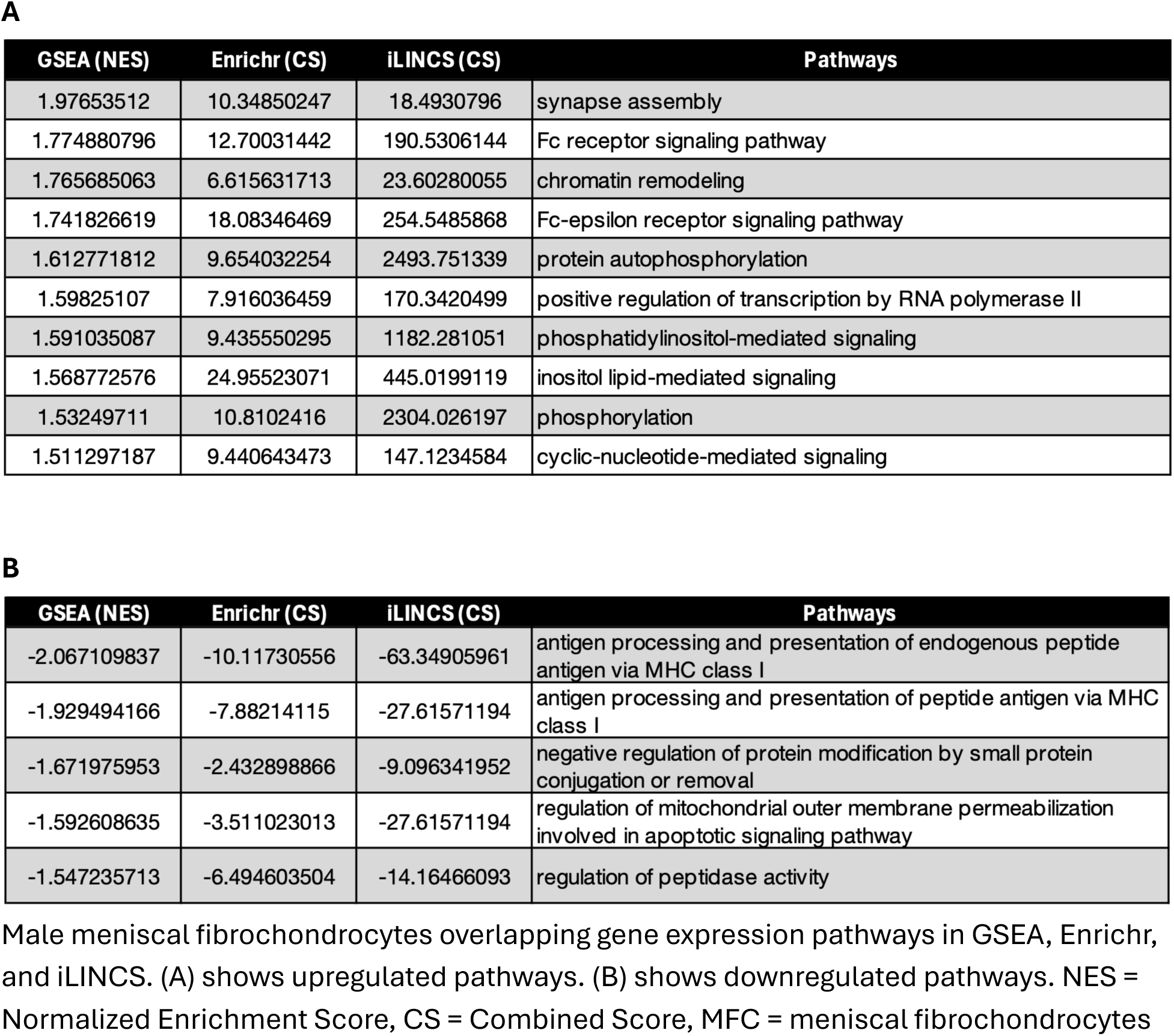
Male MFCs Overlapping Pathways in GSEA, Enrichr, and iLINCS.

**A**
| GSEA (NES) | Enrichr (CS) | iLINCS (CS) | Pathways |
| --- | --- | --- | --- |
| 1.97653512 | 10.34850247 | 18.4930796 | synapse assembly |
| 1.774880796 | 12.70031442 | 190.5306144 | Fc receptor signaling pathway |
| 1.765685063 | 6.615631713 | 23.60280055 | chromatin remodeling |
| 1.741826619 | 18.08346469 | 254.5485868 | Fc-epsilon receptor signaling pathway |
| 1.612771812 | 9.654032254 | 2493.751339 | protein autophosphorylation |
| 1.59825107 | 7.916036459 | 170.3420499 | positive regulation of transcription by RNA polymerase II |
| 1.591035087 | 9.435550295 | 1182.281051 | phosphatidylinositol-mediated signaling |
| 1.568772576 | 24.95523071 | 445.0199119 | inositol lipid-mediated signaling |
| 1.53249711 | 10.8102416 | 2304.026197 | phosphorylation |
| 1.511297187 | 9.440643473 | 147.1234584 | cyclic-nucleotide-mediated signaling |

**B**
| GSEA (NES) | Enrichr (CS) | iLINCS (CS) | Pathways |
| --- | --- | --- | --- |
| -2.067109837 | -10.11730556 | -63.34905961 | antigen processing and presentation of endogenous peptide antigen via MHC class I |
| -1.929494166 | -7.88214115 | -27.61571194 | antigen processing and presentation of peptide antigen via MHC class I |
| -1.671975953 | -2.432898866 | -9.096341952 | negative regulation of protein modification by small protein conjugation or removal |
| -1.592608635 | -3.511023013 | -27.61571194 | regulation of mitochondrial outer membrane permeabilization involved in apoptotic signaling pathway |
| -1.547235713 | -6.494603504 | -14.16466093 | regulation of peptidase activity |

Male meniscal fibrochondrocytes overlapping gene expression pathways in GSEA, Enrichr, and iLINCS. (A) shows upregulated pathways. (B) shows downregulated pathways. NES = Normalized Enrichment Score, CS = Combined Score, MFC = meniscal fibrochondrocytes

### Comparative Analysis and Sex-Specific Differences

The integrated analysis of all three methods demonstrated that female MFCs predominantly activated proliferative and DNA repair mechanisms while suppressing inflammatory and ECM-degrading processes (Table 1A,B). In contrast, male MFCs shifted towards enhancing developmental and signaling pathways, with concurrent repression of immune functions and peptidase activity (Table 2A,B).

### Leading Edge Gene Analysis

#### Leading Edge Gene Analysis in Female MFCs

The leading-edge gene analysis for female MFCs treated with 17β-estradiol (E2) identified significant genes driving pathway enrichments. For pathways involved in positive regulation of the cell cycle, key leading-edge genes included RGCC (log2FC = 0.525, p=0.001), CDCA8 (log2FC= 0.269, p<0.001), CDC45 (log2FC = 0.302, p<0.001), and CDC6 (log2FC = 0.256, p=0.006). In the DNA replication pathway, DNA2 (log2FC = 0.290, p=0.002) and CDT1 (log2FC = 0.284, p=0.002) were prominently involved (Table 3A). Conversely, the downregulated immune response pathway prominently featured MYD88 (log2FC = -0.179, p=0.004), while the suppression of metalloendopeptidase activity was notably driven by the downregulation of MME (log2FC = -0.397, p<0.001) (Table 3B). These genes represent the core drivers underpinning the significant biological changes observed following E2 treatment in female MFCs.

**Table 3:** Female MFCs Leading Edge Genes.

**A**
| Symbol | Name | N | log2FoldChange | pvalue |
| --- | --- | --- | --- | --- |
| RGCC | regulator of cell cycle | 21 | 0.525 | 0.001 |
| MCM8 | minichromosome maintenance 8 homologous recombination repair factor | 35 | 0.346 | < 0.001 |
| CDC45 | cell division cycle 45 | 28 | 0.302 | < 0.001 |
| DNA2 | DNA replication helicase/nuclease 2 | 43 | 0.29 | 0.002 |
| CDT1 | chromatin licensing and DNA replication factor 1 | 33 | 0.284 | 0.002 |
| BRCA1 | BRCA1 DNA repair associated | 41 | 0.278 | 0.002 |
| CDCA8 | cell division cycle associated 8 | 40 | 0.269 | < 0.001 |
| CDC6 | cell division cycle 6 | 36 | 0.256 | 0.006 |
| TTK | TTK protein kinase | 51 | 0.256 | 0.001 |
| RAD51 | RAD51 recombinase | 63 | 0.238 | 0.008 |
| BUB1B | BUB1 mitotic checkpoint serine/threonine kinase B | 44 | 0.184 | 0.007 |
| CDK2 | cyclin dependent kinase 2 | 37 | 0.169 | 0.01 |

**B**
| Symbol | Name | N | log2FoldChange | pvalue |
| --- | --- | --- | --- | --- |
| LBP | lipopolysaccharide binding protein | 30 | -1.468 | < 0.001 |
| MMP13 | matrix metalloproteinase 13 | 12 | -1.389 | 0.095 |
| MMP1 | matrix metalloproteinase 1 | 10 | -0.636 | 0.002 |
| IL12A | interleukin 12A | 41 | -0.528 | 0.002 |
| CXCL12 | C-X-C motif chemokine ligand 12 | 33 | -0.428 | < 0.001 |
| MME | membrane metalloendopeptidase | 14 | -0.397 | < 0.001 |
| MYD88 | MYD88 innate immune signal transduction adaptor | 25 | -0.179 | 0.004 |
| MMP11 | matrix metalloproteinase 11 | 6 | -0.154 | 0.463 |

Female meniscal fibrochondrocytes Leading Edge Genes in GSEA. (A) shows upregulated Leading Edge Genes. (B) shows downregulated Leading Edge Genes. Leading-edge genes come from the ranked GSEA list and can include genes above the $p < 0.02$ significance cutoff. MFC = meniscal fibrochondrocytes, N = number of significantly enriched GO gene sets in which the gene appears as a leading-edge member

### Leading Edge Gene Analysis in Male MFCs

The leading-edge gene analysis for male MFCs treated with 17β-estradiol (E2) highlighted key genes contributing to significant pathway enrichments. For the “synapse assembly” pathway, critical genes included PLXNA3 (log2FC = 0.250, p<0.001), SHANK3 (log2FC = 0.582, p=0.002), NOTCH1 (log2FC = 0.308, p=0.002), and LRRC4 (log2FC = 0.739, p=0.01). Additionally, upregulated chromatin remodeling pathways were primarily driven by ERCC6 (log2FC = 0.356, p<0.001), SUPT16H (log2FC = 0.136, p=0.02), BICRA (log2FC = 0.277, p=0.032), and BICRAL (log2FC = 0.347, p<0.001) (Table 4A). Conversely, the downregulated “antigen processing via MHC class I” pathway prominently involved HLA-E (log2FC = -1.266, p<0.001), HLA-A (log2FC = -0.279, p=0.016), and HLA-B (log2FC = -0.382, p<0.001). Additionally, the “regulation of peptidase activity” pathway included prominently downregulated genes such as TIMP1 (log2FC = -0.333, p<0.001), SPINT2 (log2FC = -0.412, p=0.002), SPCS1 (log2FC = -0.182, p=0.001), SPCS2 (log2FC = -0.203, p<0.001), SEC11A (log2FC = -0.151, p=0.002), SEC11C (log2FC = - 0.268, p=0.021), and PMPCB (log2FC = -0.099, p=0.014) (Table 4B). These genes represent critical contributors to the distinct biological responses observed in male MFCs following E2 treatment.

**Table 4:** Male MFCs Leading Edge Genes.

**A**
| Symbol | Name | N | log2FoldChange | pvalue |
| --- | --- | --- | --- | --- |
| SRCAP | Snf2 related CREBBP activator protein | 21 | 0.943 | < 0.001 |
| LRRC4 | leucine rich repeat containing 4 | 10 | 0.739 | 0.01 |
| SHANK3 | SH3 and multiple ankyrin repeat domains 3 | 47 | 0.582 | 0.002 |
| ERCC6 | ERCC excision repair 6, chromatin remodeling factor | 27 | 0.356 | < 0.001 |
| BICRAL | BICRA like chromatin remodeling complex associated protein | 1 | 0.347 | < 0.001 |
| NOTCH1 | notch receptor 1 | 34 | 0.308 | 0.002 |
| BICRA | BRD4 interacting chromatin remodeling complex associated protein | 1 | 0.277 | 0.032 |
| PLXNA3 | plexin A3 | 37 | 0.25 | < 0.001 |
| SUPT16H | SPT16 homolog, facilitates chromatin remodeling subunit | 2 | 0.136 | 0.02 |

**B**
| Symbol | Name | N | log2FoldChange | pvalue |
| --- | --- | --- | --- | --- |
| IFI6 | interferon alpha inducible protein 6 | 24 | -1.689 | 0.008 |
| OAS1 | 2'-5'-oligoadenylate synthetase 1 | 63 | -1.365 | 0.005 |
| HLA-E | major histocompatibility complex, class I, E | 85 | -1.266 | < 0.001 |
| SPINT2 | serine peptidase inhibitor, Kunitz type 2 | 2 | -0.412 | 0.002 |
| NDUFA3 | NADH:ubiquinone oxidoreductase subunit A3 | 65 | -0.396 | 0.001 |
| HLA-B | major histocompatibility complex, class I, B | 50 | -0.382 | < 0.001 |
| TIMP1 | TIMP metallopeptidase inhibitor 1 | 13 | -0.333 | < 0.001 |
| CST3 | cystatin C | 12 | -0.307 | < 0.001 |
| HLA-A | major histocompatibility complex, class I, A | 78 | -0.279 | 0.016 |
| SEC11C | SEC11 homolog C, signal peptidase complex subunit | 4 | -0.268 | 0.021 |
| ATP5F1D | ATP synthase F1 subunit delta | 73 | -0.268 | 0.002 |
| SPCS2 | signal peptidase complex subunit 2 | 7 | -0.203 | < 0.001 |
| SPCS1 | signal peptidase complex subunit 1 | 9 | -0.182 | 0.001 |
| SERPING1 | serpin family G member 1 | 7 | -0.16 | < 0.001 |
| SEC11A | SEC11 homolog A, signal peptidase complex subunit | 3 | -0.151 | 0.002 |
| PMPCB | peptidase, mitochondrial processing subunit beta | 4 | -0.099 | 0.014 |

Male meniscal fibrochondrocytes Leading Edge Genes in GSEA. (A) shows upregulated Leading Edge Genes. (B) shows downregulated Leading Edge Genes. Leading-edge genes come from the ranked GSEA list and can include genes above the $p < 0.02$ significance cutoff. MFC = meniscal fibrochondrocytes, N = number of significantly enriched GO gene sets in which the gene appears as a leading-edge member

## Discussion

This study provides novel insights into the sex-specific transcriptomic responses of human meniscal fibrochondrocytes (MFCs) to 17β-estradiol (E2) treatment, revealing fundamentally divergent biological programs between female and male cells. By reanalyzing RNA sequencing data from dataset GSE199087 and integrating three independent pathway analysis platforms—GSEA, Enrichr, and iLINCS—we demonstrate that E2 activates proliferative and DNA repair pathways in female MFCs while simultaneously suppressing inflammatory and matrix-degrading processes. In contrast, male MFCs respond to E2 through chromatin remodeling and developmental signaling pathways with concurrent repression of immune functions. These divergent responses carry significant implications for understanding the well-documented sexual dimorphism in osteoarthritis (OA) prevalence and severity, where females carry a disproportionate disease burden, particularly after menopause (*15, 16*).

### E2-Mediated Protection of Female MFCs: A Dual Mechanism

In female MFCs, E2 treatment produced a coordinated upregulation of genes governing cell cycle progression and DNA repair, including RGCC (log^2^FC = 0.525, p = 0.001), MCM8, CDC45, BRCA1, RAD51, BUB1B, and CDK2 (Table 3A). Three representative overlapping pathways—DNA-templated DNA replication (GSEA NES = 2.15), DNA replication (GSEA NES = 2.14), and positive regulation of cell cycle process (GSEA NES = 1.78, Enrichr CS = 304.79)—indicate a robust proliferative and genomic maintenance response. Concurrently, E2 drove significant downregulation of genes associated with inflammation and ECM catabolism, including MMP1 (log^2^FC = −0.636, p = 0.002), LBP, IL12A, CXCL12, MYD88 (log^2^FC = −0.179, p = 0.004), and MME (log^2^FC = −0.397, p < 0.001) (Table 3B). The table additionally lists non-significant members such as MMP11 (log2FC = −0.154, p = 0.463), included to show which catabolic genes did and did not respond to E2. Downregulated pathways converged on endopeptidase activity, fibroblast growth factor receptor signaling, regulation of defense response, and positive regulation of cell death (Table 1B).

This dual mechanism—enhancing repair while suppressing degradation—is consistent with the established chondroprotective actions of estrogen in articular cartilage. Claassen et al. demonstrated that 17β-estradiol suppresses mRNA levels of MMP-3 and MMP-13 in articular chondrocytes from female patients cultured in a three-dimensional alginate system, with MMP-1 reduced at higher doses (*2*). Similarly, Lee et al. reported that E2 reduces MMP-1 secretion in OA chondrocytes, particularly in the absence of high-level cytokine stimulation (*3*). The downregulation of MMP1 in our female MFC dataset, alongside a pronounced but non-significant downward trend in MMP13 (log2FC = −1.389, p = 0.095), extends these findings from articular cartilage to meniscal fibrocartilage, suggesting that estrogen’s anti-catabolic effects may operate across multiple cartilaginous tissues of the knee. Beyond direct MMP suppression, E2 has been shown to protect chondrocytes through additional mechanisms, including promotion of autophagy and mitophagy via the SIRT1-mediated AMPK/mTOR signaling pathway (*17*) and regulation of MMP-13 expression through microRNA-140 (*4*). Whether these alternative protective mechanisms contribute to the transcriptomic changes observed in female MFCs warrants further investigation.

### Distinct Transcriptomic Response in Male MFCs

Male MFCs displayed a markedly different transcriptomic response to E2. Upregulated genes were enriched for synapse assembly, chromatin remodeling, and receptor signaling, with prominent leading edge genes including SRCAP (log^2^FC = 0.943, p < 0.001), SHANK3 (log^2^FC = 0.582, p = 0.002), ERCC6 (log^2^FC = 0.356, p < 0.001), NOTCH1 (log^2^FC = 0.308, p = 0.002), BICRAL, BICRA, and SUPT16H (Table 4A). Three representative overlapping upregulated pathways—synapse assembly (GSEA NES = 1.98), chromatin remodeling (GSEA NES = 1.77), and Fc receptor signaling (GSEA NES = 1.77)—indicate that male MFCs prioritize epigenetic and developmental programs rather than the proliferative and repair-oriented response seen in female cells.

The upregulation of NOTCH1 in male MFCs is of particular interest given the central role of Notch signaling in cartilage pathobiology. NOTCH1 is the most abundantly expressed Notch receptor in articular cartilage, and its expression is elevated in OA, where it drives chondrocyte hypertrophy and dedifferentiation (*18, 19*). Hosaka et al. showed that tissue-specific inactivation of the Notch transcriptional effector RBPjκ in adult articular cartilage confers resistance to OA development in mouse knee joints, while Notch intracellular domain overexpression stimulates endochondral ossification through induction of Hes1 (*18*). Minguzzi et al. further demonstrated that silencing NOTCH1 in human OA chondrocytes reduces proliferation, decreases expression of hypertrophy markers RUNX2 and MMP-13, and improves viability in 3D culture (*19*). In this context, the E2-mediated upregulation of NOTCH1 in male MFCs may represent a potentially deleterious signal that could promote hypertrophic differentiation rather than tissue maintenance.

On the downregulated side, male MFCs showed striking repression of MHC class I antigen presentation genes (HLA-A, HLA-B, HLA-E) and mitochondrial components (NDUFA3, ATP5F1D), as well as TIMP1 (log^2^FC = −0.333, p < 0.001) (Table 4B). The downregulation of TIMP1 in male cells deserves special attention, as TIMPs are endogenous inhibitors of MMPs. While female MFCs downregulated the MMPs themselves, male MFCs instead lost expression of a key MMP inhibitor—a distinction that could shift the proteolytic balance toward ECM degradation through different mechanisms. This contrast underscores that E2 modulates the MMP/TIMP axis in sex-divergent ways in meniscal tissue.

### Implications for OA Sexual Dimorphism and Menopausal Risk

The sex-specific patterns observed here may help explain the pronounced increase in OA incidence that women experience around menopause. A meta-analysis by Srikanth et al. found that males had a significantly reduced risk for prevalent knee OA compared to females (RR 0.63, 95% CI 0.53–0.75), with females tending toward more severe disease particularly after menopausal age (*15*). More recently, Segal et al. reported that women account for approximately 60% of all OA cases globally, with the sex disparity widening after age 40 (*16*). The decline in circulating estrogen at menopause has been proposed as a contributing factor to this increased susceptibility, supported by evidence that estrogen deficiency accelerates cartilage degradation and promotes inflammatory cytokine expression in joint tissues (*20, 21*).

Our transcriptomic data suggest a plausible mechanism: in the presence of E2, female MFCs maintain an actively protective phenotype characterized by cell renewal and inflammatory suppression. The withdrawal of E2 at menopause would then remove this dual protective program, potentially exposing meniscal tissue to unchecked catabolic and inflammatory processes. This interpretation aligns with some epidemiological evidence suggesting that hormone replacement therapy may offer modest protective effects against radiographic knee OA, although results across studies remain inconsistent (*22, 23*). Jung et al. reported a reduced odds ratio (0.70, 95% CI 0.50–0.99) for knee OA in postmenopausal women receiving menopausal hormone therapy in a nationwide Korean cohort (*23*), while other studies have yielded conflicting results, highlighting the complexity of estrogen’s role in joint health.

### Estrogen Receptor Expression in Meniscal Tissue

The differential response to E2 between sexes may be partially explained by variations in estrogen receptor expression across joint tissues. Wang et al. demonstrated in a mouse model that ER-α and ER-β are differentially expressed across fibrocartilaginous tissues, with relatively weak expression of both receptors in knee meniscus cells compared to temporomandibular joint and pubic symphysis cells (*24*). Although this study was conducted in mice and its applicability to human meniscal cells remains to be established, it suggests that meniscal cells may respond to E2 through pathways that differ from those in articular chondrocytes, which have been more extensively characterized. Whether sex-based differences in receptor density or subtype distribution contribute to the divergent transcriptomic responses observed in our study is an important question for future work.

### Clinical Implications

These findings carry several clinical implications. First, they provide molecular evidence that estrogen-based interventions for OA should account for sex as a biological variable, since male and female meniscal cells activate fundamentally different programs in response to E2. Second, the protective effects of E2 in female MFCs—promoting repair while suppressing catabolism—provide biological rationale for investigating estrogen-based strategies to preserve meniscal health in post-menopausal women, though any such approach must be weighed against the broader risk–benefit profile of hormone therapy. Third, the identification of specific pathway alterations in each sex offers potential pharmacological targets. For example, the pronounced downregulation of MMP1 and MYD88 in female cells suggests that targeted anti-catabolic therapies could complement estrogen’s natural protective effects, while the upregulation of NOTCH1 in male cells points to the Notch pathway as a potential axis of intervention that has already shown therapeutic promise in preclinical OA models (*18, 25*). Pan et al. have previously highlighted that OA in human knees exhibits sex-specific molecular characteristics, reinforcing the need for sex-stratified approaches to both research and treatment (*6*).

### Limitations

Several limitations warrant consideration. Most importantly, our analysis was based on MFCs derived from one male and one female cadaveric donor (both aged 47 years), which substantially limits the generalizability of our findings. While triplicate experiments and pulsed dosing enhance the reliability of the transcriptomic data within this dataset, findings from single-donor analyses cannot capture the range of individual variation in estrogen responsiveness that exists across populations. Second, the in vitro monolayer culture conditions do not recapitulate the complex in vivo environment of the knee joint, where MFCs experience mechanical loading, interact with other cell types, and are embedded within an organized extracellular matrix. Third, the pulsed dosing regimen (1 hour E2 followed by 23 hours E2-free, repeated over 72 hours), while designed to mimic cyclical estrogen exposure, may not precisely replicate physiological hormonal dynamics. Fourth, our analysis is based on transcriptomic changes alone; protein-level validation of key differentially expressed genes would be needed to confirm functional significance. Finally, as a secondary analysis of a publicly available dataset (GSE199087), our study is constrained by the experimental parameters of the original investigators.

### Future Directions

Future studies should investigate E2 effects on MFCs from a larger, more diverse cohort of donors to assess reproducibility and capture individual variation. Protein-level validation of key findings—particularly MMP1, MMP13, MYD88, NOTCH1, and TIMP1—would strengthen the transcriptomic observations. The use of more physiologically relevant models, such as 3D culture systems or ex vivo meniscal explants, would better approximate the in vivo meniscal environment, and the plasticity of MFCs in such systems has been previously documented (*5*). Investigating estrogen receptor subtype expression (ER-α vs. ER-β) in human meniscal tissue and its relationship to the observed sex-specific responses would help clarify the signaling mechanisms involved. Furthermore, examining the effects of other sex hormones—including progesterone and testosterone—on MFC gene expression would provide a more complete picture of hormonal influences on meniscal biology. Longitudinal studies correlating serum estrogen levels, meniscal integrity, and OA progression in human subjects would help validate these in vitro observations clinically.

## Conclusion

Our study demonstrates that 17β-estradiol elicits sex-specific transcriptomic responses in human meniscal fibrochondrocytes. Female cells activate proliferative, DNA repair, and cell cycle pathways while suppressing inflammatory and ECM-degrading processes—a dual protective program that is lost with estrogen decline at menopause. Male cells instead engage chromatin remodeling and developmental signaling pathways, including NOTCH1 upregulation, with concurrent repression of immune functions and TIMP1 expression. These findings extend our understanding of estrogen’s chondroprotective effects from articular cartilage to meniscal tissue and provide a molecular framework for the sexual dimorphism observed in OA. The identification of sex-specific pathway responses supports the development of personalized, sex-informed therapeutic strategies for meniscal preservation and osteoarthritis management.

